# Combination therapy with oral antiviral and anti-inflammatory drugs improves the efficacy of delayed treatment in severe COVID-19

**DOI:** 10.1101/2023.06.20.545832

**Authors:** Michihito Sasaki, Tatsuki Sugi, Shun Iida, Yuichiro Hirata, Shinji Kusakabe, Kei Konishi, Yukari Itakura, Koshiro Tabata, Mai Kishimoto, Hiroko Kobayashi, Takuma Ariizumi, Kittiya Intaruck, Haruaki Nobori, Shinsuke Toba, Akihiko Sato, Keita Matsuno, Junya Yamagishi, Tadaki Suzuki, William W. Hall, Yasuko Orba, Hirofumi Sawa

**Affiliations:** Division of Molecular Pathobiology, International Institute for Zoonosis Control, Hokkaido University, Sapporo, Japan; Institute for Vaccine Research and Development (HU-IVReD), Hokkaido University, Sapporo, Japan; Division of Collaboration and Education, International Institute for Zoonosis Control, Hokkaido University, Sapporo, Japan; International Collaboration Unit, International Institute for Zoonosis Control, Hokkaido University, Sapporo, Japan; Department of Pathology, National Institute of Infectious Diseases, Tokyo, Japan; Division of Anti-Virus Drug Research, International Institute for Zoonosis Control, Hokkaido University, Sapporo, Japan; Drug Discovery & Disease Research Laboratory, Shionogi & Co., Ltd., Osaka, Japan; Division of Risk Analysis and Management, International Institute for Zoonosis Control, Hokkaido University, Sapporo, Japan; One Health Research Center, Hokkaido University, Sapporo, Japan; National Virus Reference Laboratory, School of Medicine, University College of Dublin, Ireland; Global Virus Network, Baltimore, Maryland, USA

## Abstract

Pulmonary infection with SARS-CoV-2 stimulates host immune responses and can also result in the progression of dysregulated and critical inflammation. Throughout the pandemic, the management and treatment of COVID-19 has been continuously updated with a range of antiviral drugs and immunomodulators. Monotherapy with oral antivirals has proven to be effective in the treatment of COVID-19. However, the treatment should be initiated in the early stages of infection to ensure beneficial therapeutic outcomes, and there is still room for further consideration on therapeutic strategies using antivirals. Here, we show that the oral antiviral ensitrelvir combined with the anti-inflammatory corticosteroid methylprednisolone has higher therapeutic effects and better outcomes in a delayed dosing model of SARS-CoV-2 infected hamsters compared to the monotherapy with ensitrelvir or methylprednisolone alone. Combination therapy with these drugs improved respiratory conditions and the development of pneumonia in hamsters even when the treatment was started after 2 days post infection. The combination therapy led to a differential histological and transcriptomic pattern in comparison to either of the monotherapies, with reduced lung damage and down-regulated expressions of genes involved in inflammatory response. Furthermore, we found that the combination treatment is effective in infection with both highly pathogenic delta and circulating omicron variants. Our results demonstrate the advantage of combination therapy with antiviral and corticosteroid drugs in COVID-19 treatment. Since both drugs are available as oral medications, this combination therapy could provide a clinical and potent therapeutic option for COVID-19.

## Introduction

Coronavirus disease 2019 (COVID-19), caused by infection with severe acute respiratory syndrome coronavirus 2 (SARS-CoV-2), is linked to mild-to-severe respiratory distress, pneumonia, and death. Bronchial and alveolar epithelial cells are the major target cells for SARS-CoV-2 infection, and these cells trigger host innate immune and inflammatory responses in the lungs (Lamers & Haagmans, 2022). Following viral replication, interferons and cytokines are produced and released from infected and bystander epithelial cells and immune cells, leading to the activation and pulmonary infiltration of immune cells including macrophages and neutrophils (Hsu *et al*, 2022). Although a proper immune response orchestrates the antiviral state and attenuates tissue damages by viral infection, severe COVID-19 shows an imbalanced and hyperactive host immune response characterized by defects in the type I interferon response and the dysregulated release of pro-inflammatory cytokines, resulting in deleterious inflammation (Diamond & Kanneganti, 2022; Merad *et al*, 2022). Overall, the inflammatory response to SARS-CoV-2 infection plays a pivotal role in the pathogenesis of severe COVID-19.

Several medications have been employed for COVID-19 therapy (COVID-19 Treatment Guidelines Panel. 2023). In the early phase of infection, oral antiviral agents such as nirmatrelvir and molnupiravir are recommended for treating non-hospitalized patients to reduce the viral load and the risk of COVID-19-related hospitalization (Hammond *et al*, 2022; Jayk Bernal *et al*, 2021). In addition, ensitrelvir (ETV) reduces the time to resolution of COVID-19 symptoms, and it is currently approved as an oral antiviral medication for COVID-19 in Japan (Mukae *et al*, 2023). For therapeutic efficacy, it is disable that these oral antiviral medications are initiated as soon as possible after the onset of symptoms (COVID-19 Treatment Guidelines Panel. 2023). In the later stages of infection, immunomodulators are expected to circumvent tissue damage induced by the inflammatory response observed exclusively in severe COVID-19. In such a situation, dexamethasone, an anti-inflammatory corticosteroid, could reduce the mortality rate in hospitalized patients with COVID-19 requiring supplemental oxygen (Horby *et al*, 2021). Methylprednisolone (mPSL) is also used as an alternative corticosteroid for the same purpose, while the evidence of the efficacy of mPSL in severe COVID-19 remains insufficient because of the small sample sizes in clinical trials (Hong *et al*, 2023). Because corticosteroids cause immunosuppression and increase viral loads, this treatment is harmful rather than beneficial in patients with mild-to-moderate COVID-19 (Covello *et al*, 2023; Crothers *et al*, 2022). Meanwhile, combination therapy with the intravenous antiviral drug remdesivir and immunomodulators produced better outcomes than treatment with remdesivir or immunomodulator alone in severe diseases (Kalil *et al*, 2021; Marrone *et al*, 2022). However, little is known about the therapeutic impact of combination therapy with oral antiviral agents and immunomodulators in both mild-to-moderate and severe COVID-19.

Syrian hamsters are highly susceptible to infection with SARS-CoV-2, and they are widely used in *in vivo* studies of COVID-19. To date, the hamster has proven to be the most reliable animal model for studying COVID-19 pathophysiology and therapeutics. In preclinical studies, oral antiviral agents reduced viral loads and improved lung pathology in recipient hamsters infected with SARS-CoV-2 (Abdelnabi *et al*, 2022; Rosenke *et al*, 2021; Uraki *et al*, 2022b). Recently, we demonstrated that post-exposure treatment with ETV in SARS-CoV-2-infected hamsters resulted in reduced viral replication and alleviated symptoms in hamsters (Sasaki *et al*, 2023). Treatment with corticosteroids also improved pathology but increased viral loads and delayed clearance of virus in the lungs of hamsters (Wyler *et al*, 2022; Ye *et al*, 2021; Yuan *et al*, 2022a). Moreover, two studies examined the combinations of remdesivir plus mPSL (Ye *et al*., 2021) and neutralizing monoclonal antibody plus dexamethasone (Wyler *et al*., 2022), observing favorable therapeutic effects compared to those of monotherapy.

Considering the relationship between virus replication and host inflammatory response in lungs, it is desirable to initiate treatment with oral antiviral drugs early after the onset of COVID-19. However, treatment in clinical settings is sometimes initiated in the context of extensive SARS-CoV-2 replication in the respiratory tracts of patients. In this study, we investigated the influence of delaying oral antiviral ETV treatment on therapeutic efficacy using a SARS-CoV-2-infected hamster model. We then evaluated and characterized the therapeutic effect of oral combination treatment with ETV and mPSL in a delayed dosing model using histopathological and transcriptome analyses.

## Results

### Delayed treatment attenuates the therapeutic effect of ETV in hamsters

To examine the influence of delaying antiviral administration on COVID-19 treatment, hamsters were intranasally infected with SARS-CoV-2 and then orally treated for 5 days with ETV using three different starting times: immediately after inoculation to 4 days post-infection (dpi) [ETV (0 dpi)], from 1 to 5 dpi [ETV (1 dpi)], and from 2 to 6 dpi [ETV (2 dpi); Fig. 1A]. Hamsters treated with methyl cellulose 400 (vehicle) were included as an untreated control group in this study. We used the SARS-CoV-2 delta variant (lineage AY.122), which is a highly pathogenic variant that causes severe pneumonia in hamsters (Halfmann *et al*, 2022; Saito *et al*, 2022; Yuan *et al*, 2022b). Vehicle-treated hamsters exhibited body weight loss up to 6 dpi (Fig. 1B). Oral ETV alleviated the reduction of body weight associated with the infection, whereas delayed treatment had limited ameliorating effects on body weight loss. The respiratory condition of hamsters at 4 dpi was assessed by measuring the enhanced pause (Penh) and ratio of peak expiratory flow (Rpef) using whole-body plethysmography (Fig. 1C, D). ETV treatment inhibited the increase in Penh and decrease in Rpef associated with SARS-CoV-2 infection, indicating improvement in the respiratory condition, but this effect was limited in the delayed-treatment groups. Lung tissues were harvested from a subset of hamsters at 4 dpi in each group. Early treatment with ETV resulted in a larger decrease in viral loads in the lungs, but the viral loads were also decreased by 1–2 orders of magnitude even when treatment was started after 2 dpi (Fig. 1E, F). On gross examination, focal congestion and hemorrhage were observed in the lungs of hamsters in the vehicle and ETV (2 dpi) groups, but such findings were limited in hamsters in the ETV (0 dpi) and ETV (1 dpi) groups (Fig. 1G). The cytokines IL-6, IL-10 and the chemokines CCL2/MCP-1, CXCL10/IP-10 are associated with severe COVID-19, and can serve as biological markers for COVID-19 severity (Hsu *et al*., 2022; Lamers & Haagmans, 2022). The gene expression of these cytokines and chemokines was induced in the lungs of infected hamsters at 4 dpi, and this upregulation was strongly inhibited in the ETV (0 dpi) and ETV (1 dpi) groups but only slightly inhibited in the ETV (2 dpi) group (Fig. 1H–K). These results indicate that ETV treatment reduces the viral load and ameliorates both the inflammatory state in the lungs and the clinical aspects of COVID-19, but the therapeutic effect is mitigated when ETV treatment is delayed after infection.

**Fig. 1.**
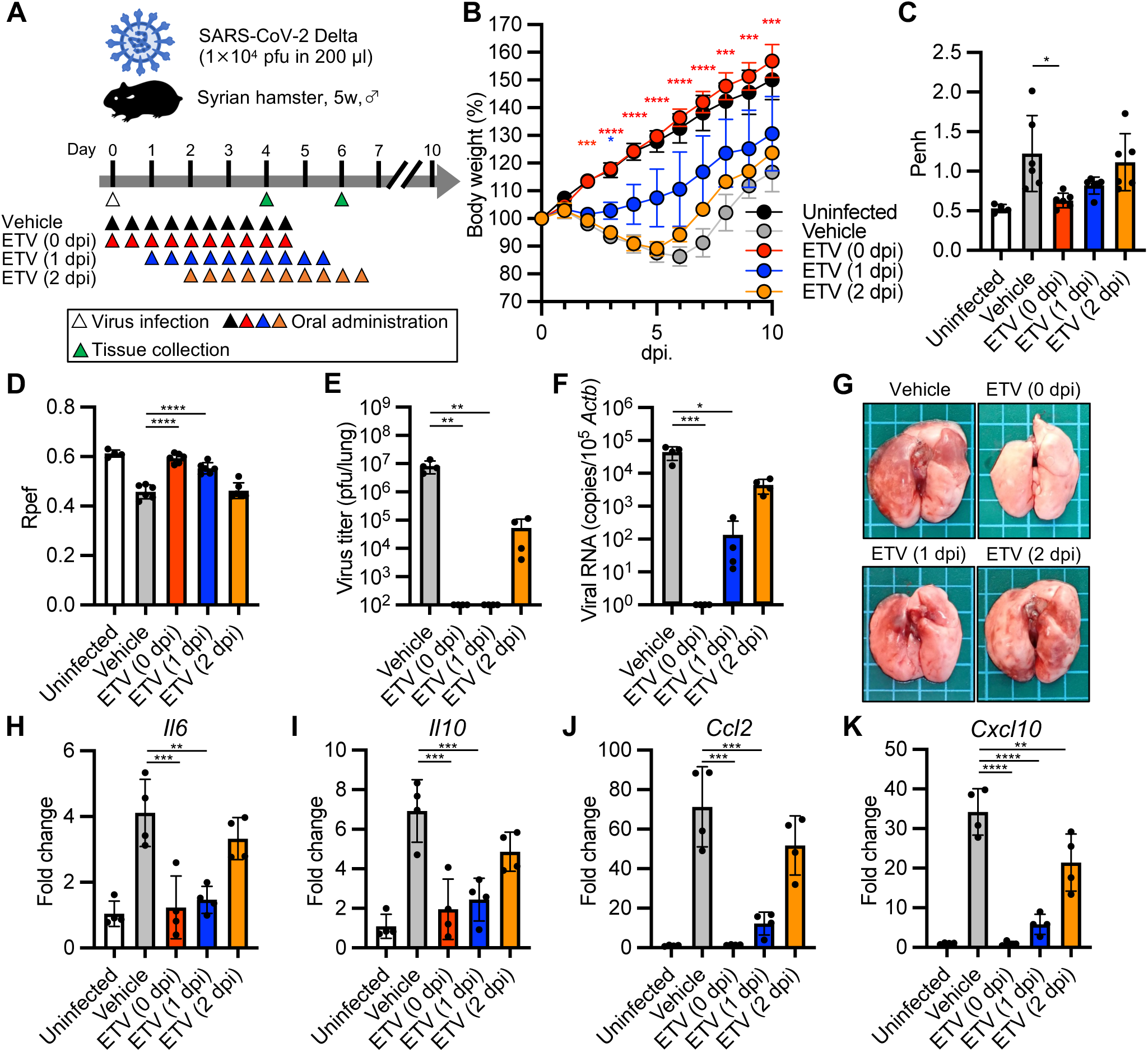
Differences in outcomes between immediate and delayed treatment with the antiviral drug ensitrelvir (ETV) in hamsters infected with SARS-CoV-2. **A**, Schematic representation of the experimental design for antiviral treatment in the hamster model. Hamsters were intranasally inoculated with 1 × 10^4^ pfu of the SARS-CoV-2 delta variant. The hamsters were orally treated with ensitrelvir (ETV, 200 mg/kg) or vehicle alone b.i.d. for 5 days starting at the time of infection [ETV (0 dpi) and vehicle], 1 day post-infection (dpi) [ETV (1 dpi)], or 2 dpi [ETV (2 dpi)]. A group of hamsters was sacrificed at 4 dpi for tissue collection. Another subset of hamsters was monitored for 10 dpi for body weight changes. **B**, Body weight changes in uninfected and SARS-CoV-2-infected hamsters treated with ETV or vehicle (n = 4 for each group). **C, D**, Enhanced pause (Penh) (C) and ratio of peak expiratory flow (Rpef) (D) in hamsters at 4 dpi were measured by whole-body plethysmography (n = 4 for the uninfected group, n = 6 for the other groups). **E**, Virus titers in the lungs of hamsters at 4 dpi were determined by the plaque assay. **F**, Viral RNA copies in the lungs of hamsters at 4 dpi were quantified by quantitative RT-PCR (qRT-PCR) and normalized to those of *Actb*. **G**, Gross observation of lungs at 4 dpi. Side length of squares, 5 mm. **H**–**K**, Relative gene expression of *Il6* (H), *Il10* (I), *Ccl2* (J), and *Cxcl10* (K) in infected hamster lungs at 4 dpi compared to that in the lungs of uninfected hamsters were examined using qRT-PCR. Data were normalized to those of *Actb*. The values shown are mean ± SD with each dot representing an individual animal. \**p*<0.05, \*\**p*<0.01, \*\*\**p*<0.001, \*\*\*\**p*<0.0001 by two-way ANOVA with Dunnett’s test (B), one-way ANOVA with Tukey’s test (C, D, H–K) and Kruskal-Wallis test with Dunn’s test (E, F).

### Combination therapy with ETV and mPSL accelerates the recovery from severe COVID-19

Previous studies demonstrated that the administration of anti-inflammatory corticosteroids (dexamethasone and mPSL) ameliorates pneumonia temporally but increases the viral load in hamsters infected with SARS-CoV-2 (Wyler *et al*., 2022; Ye *et al*., 2021; Yuan *et al*., 2022a). We next examined the therapeutic effect of delayed corticosteroid treatment in hamsters infected with SARS-CoV-2. Hamsters infected with the SARS-CoV-2 delta variant were orally administered with mPSL for 2 days (2–3 dpi) and/or ETV for 5 days (2–6 dpi; Fig. 2A). Treatment with mPSL alone had little impact on body weight loss caused by infection, but the hamsters receiving ETV/mPSL combination therapy showed the lowest body weight loss among all examined groups (Fig. 2B). Penh and Rpef at 4 dpi were improved by mPSL alone and in combination with ETV, whereas mPSL monotherapy aggravated the respiratory condition in hamsters at 6 dpi (Fig. 2C, D). Infectious virus and viral RNA loads in the lungs of hamsters at 4 and 6 dpi were reduced by ETV/mPSL combination therapy, but their viral loads were slightly higher than those in hamsters receiving ETV monotherapy (Fig. 2E–H). Hamsters treated with mPSL monotherapy maintained relatively high viral loads at 6 dpi compared to those in vehicle-treated hamsters, consistent with a previous study (Ye *et al*., 2021). Collectively, these results indicate that ETV/mPSL combination treatment exhibits high therapeutic activity and ameliorates the severity of COVID-19 without delaying virus clearance in a delayed dosing model of SARS-CoV-2 infected hamsters.

**Fig. 2.**
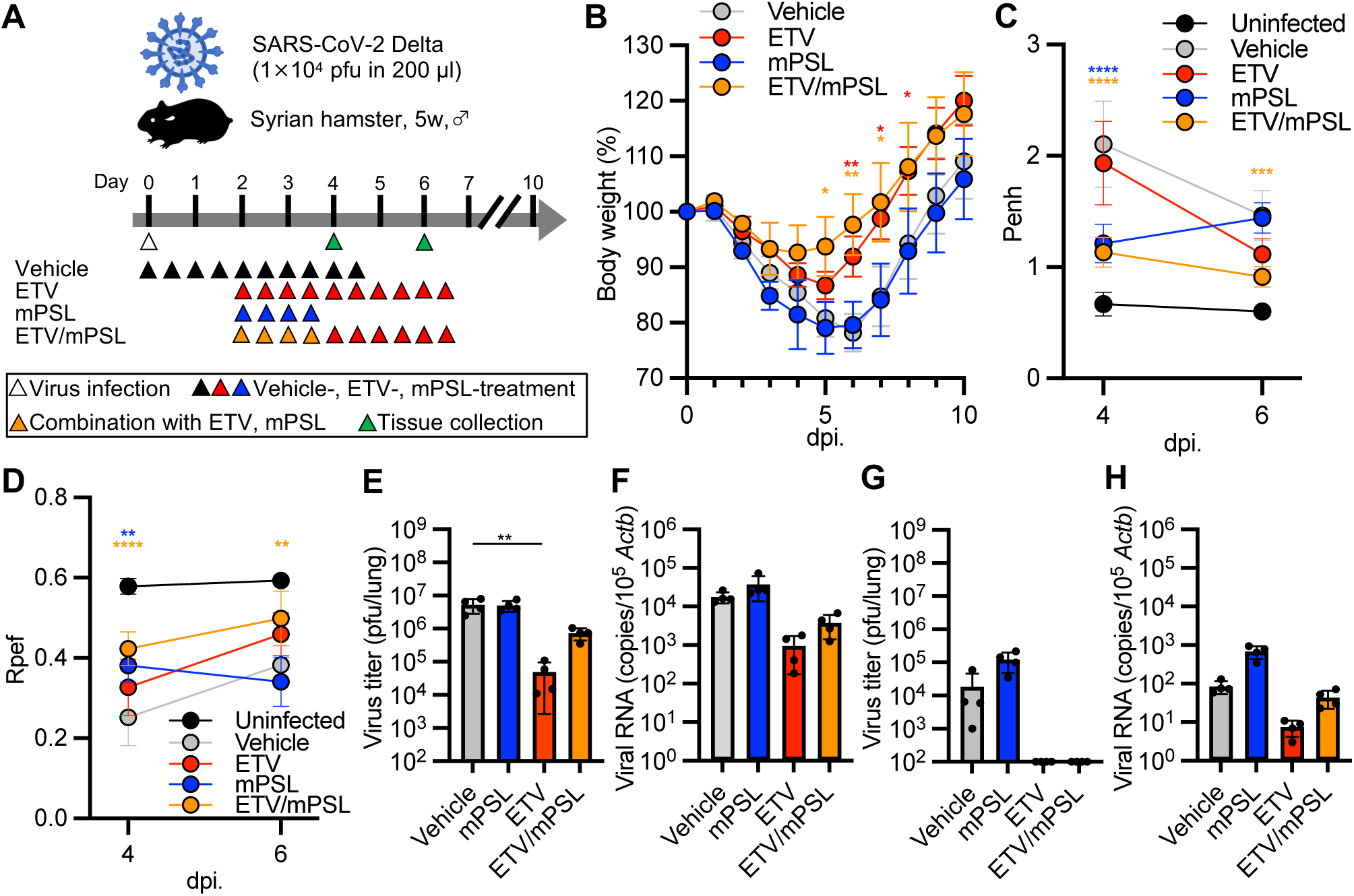
Therapeutic treatment of SARS-CoV-2 infection in hamsters with ETV and methylprednisolone (mPSL). **A**, Schematic represenatation of the experimental design for treatment in the hamster model. Hamsters inoculated with the SARS-CoV-2 delta variant were orally treated with ETV for 5 days (200 mg/kg, b.i.d., 2–6 dpi) and/or mPSL for 2 days (1 mg/kg, b.i.d., 2–3 dpi). A group of hamsters was sacrificed at 4 dpi for tissue collection. Another subset of hamsters was monitored for 10 dpi for body weight changes. **B**, Body weight changes in SARS-CoV-2-infected hamsters treated with ETV and/or mPSL (n = 4 for each group). **C, D**, Enhanced pause (Penh) (C) and ratio of peak expiratory flow (Rpef) (D) in hamsters at 4 and 6 dpi were measured by whole-body plethysmography (n = 4 for the uninfected group, n = 6 for the other groups). **E–H,** Virus titers (E, G) and viral RNA copies (F, H) in the lungs of hamsters at 4 (E, F) and 6 dpi (G, H). Viral RNA copies were normalized to those of *Actb*. The values shown are mean ± SD with each dot representing an individual animal. \**p*<0.05, \*\**p*<0.01, \*\*\**p*<0.001, \*\*\*\**p*<0.0001 by two-way ANOVA with Dunnett’s test (B–D) and Kruskal-Wallis test with Dunn’s test (E–H).

### Effects of combination therapy with ETV and mPSL on lung pathology

We examined the pathological features in hamster lungs at 4 dpi. On gross examination, focal congestion and hemorrhage were observed in the lungs of hamsters treated with ETV or mPSL alone as well as in the vehicle control (Fig. 3A). In contrast, few pathological changes were observed in the lungs of animals that received ETV/mPSL combination therapy. The histopathological severity score of pneumonia was determined on the basis of inflammation, edema, and hemorrhage as described in the Materials and Methods. Histopathological examination uncovered extensive inflammatory cell infiltration along with alveolar hemorrhage in the lungs of vehicle-treated hamsters (Fig. 3B). This severe inflammation was slightly but insignificantly reduced in the lung sections of hamsters treated with mPSL or ETV monotherapy (Fig. 3C). Meanwhile, lung sections of hamsters treated with ETV/mPSL combination therapy demonstrated mild inflammation in limited legions and significantly lower severity scores than in the vehicle-treated animals (Fig. 3B, C). Immunohistochemical and *in situ* hybridization analyses revealed that viral antigen-and viral RNA-positive cells were distributed across extended areas of the lungs of the hamsters in the vehicle and mPSL monotherapy groups, whereas a smaller number of cells in the lungs of animals in the ETV monotherapy and ETV/mPSL combination therapy groups were positive for viral RNA/antigen (Fig. 3B). Because hamsters receiving ETV/mPSL combination therapy exhibited mild symptoms with limited inflammatory responses to the infection during the monitoring period, we evaluated seroconversion in the hamsters. Hamsters treated with vehicle, ETV alone, mPSL alone, and ETV/mPSL combination therapy elicited comparable neutralizing antibody titers at 18 dpi (Fig. 3D). Taken together, ETV/mPSL combination treatment restricted both viral spread and hyperinflammation in the lungs without the change in seroconversion to SARS-CoV-2 infection, leading to more desirable therapeutic effects compared to that of ETV or mPSL monotherapy.

**Fig. 3.**
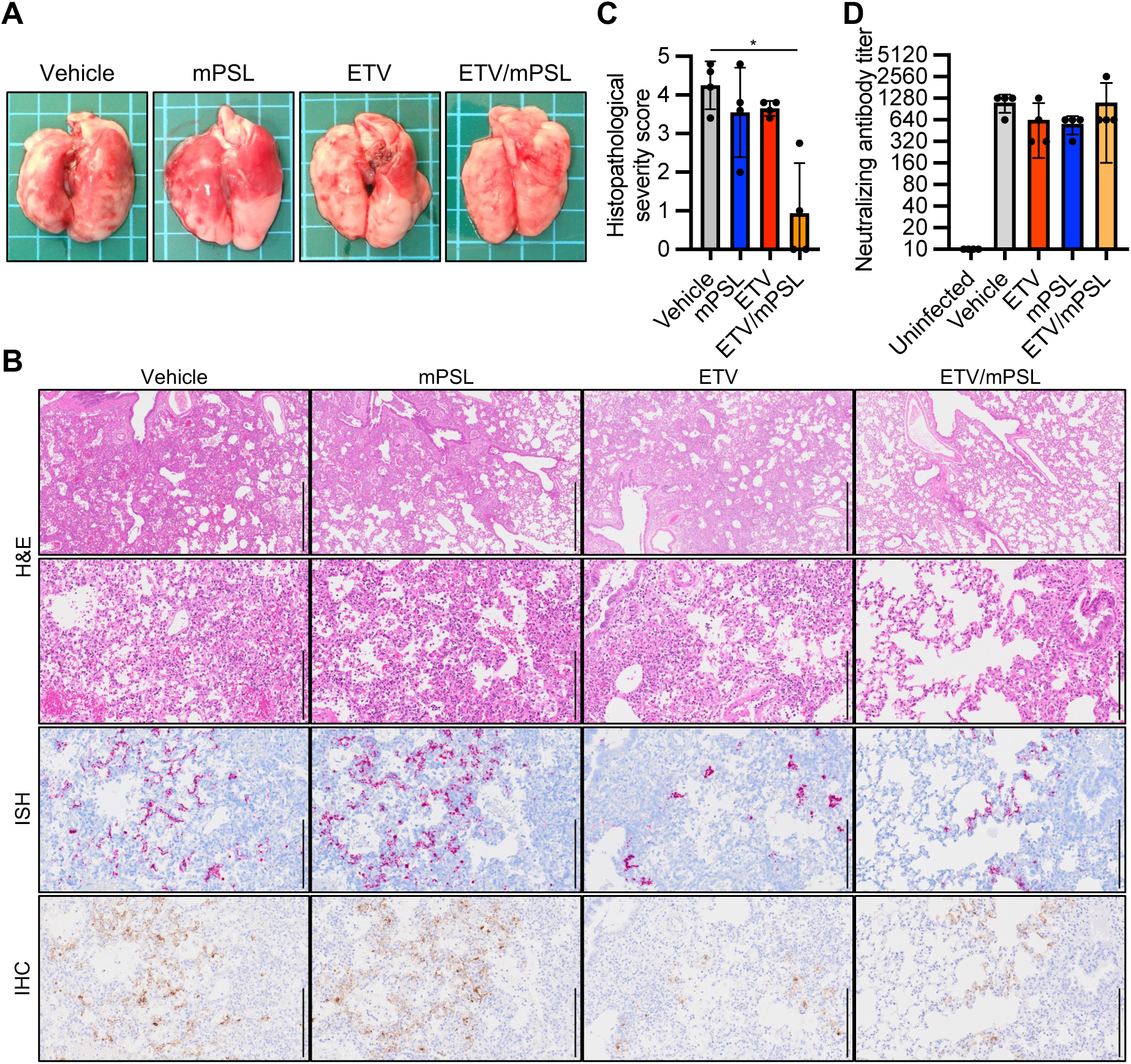
Histopathological findings in the lungs of SARS-CoV-2-infected hamsters receiving therapeutic treatment. Hamsters were infected with 1 × 10^4^ pfu of the SARS-CoV-2 delta variant and treated b.i.d. with ETV (200 mg/kg) and/or mPSL (1 mg/kg) at 2–3 dpi following the schedule shown in Fig. 2A. **A,** Gross observation of lungs from hamsters at 4 dpi. Side length of squares, 5 mm. **B,** Representative histopathological images of the lungs of hamsters at 4 dpi. Upper two panels, hematoxylin and eosin staining. The third from the top panels, *in situ* hybridization (ISH) targeting the nucleocapsid gene of SARS-CoV-2. Bottom panels, immunohistochemistry (IHC) targeting the nucleocapsid protein of SARS-CoV-2. Scale bars in the top panels, 1 mm. Scale bars in other panels, 200 μm. **C**, Histopathological severity score of pneumonia based on the percent area of alveolitis in a given section. **D**, Neutralizing antibody titers in hamster serum collected at 18 dpi. The values shown are mean ± SD with each dot representing an individual animal. \**p*<0.05 by two-way ANOVA with Kruskal-Wallis test with Dunn’s test.

### Combination therapy with ETV and mPSL controls the host inflammatory response to COVID-19

SARS-CoV-2 infection induces marked and imbalanced upregulation of various cytokines, chemokines, and interferon-stimulated genes (ISGs), which are associated with COVID-19 pneumonia and disease severity. We therefore extended our analysis of the effect of treatment on gene expression profiles. The transcriptome of hamster lungs at 4 dpi was analyzed by mRNA sequencing (mRNA-seq). Principal component analysis revealed distinct gene expression signatures between uninfected and infected (vehicle-treated) hamsters (Fig. 4A). The gene expression signatures of hamsters treated with ETV/mPSL combination therapy were separated from those of hamsters treated with vehicle, ETV, or mPSL alone (Fig. 4A). Consistent with the results of the histopathological examination, the differential expression of cell marker genes suggested a prominent inflammatory cell infiltration in the lungs of hamsters in the vehicle, ETV, and mPSL groups compared to that in the ETV/mPSL combination therapy group (Fig. S1). Pathway analysis of differentially expressed genes highlighted host innate immune and inflammatory responses to the infection, such as cytokine/chemokine signaling and cell cycle regulation, in infected hamsters compared to uninfected hamsters (Fig. 4B, left panel). In comparison to vehicle treatment, mPSL monotherapy inhibited the upregulation of some genes enriched for neutrophil degranulation and T cell receptor (TCR) signaling, whereas ETV monotherapy had little impact on these pathways (Fig. 4B, right panel and 4C). Notably, ETV/mPSL combination therapy reverted the upregulation of genes enriched for interferon and interleukin signaling in addition to neutrophil degranulation and TCR signaling by infection (Fig. 4B and 4C), indicating a comprehensive effect of treatment on the host immune response to COVID-19. We validated the expression of cytokines and chemokines in hamster lungs at 4 and 6 dpi by quantitative RT-PCR (qRT-PCR; Fig. 5A, B). We also included *Oas2* and *Irf7*, which were upregulated ISGs in our transcriptome analysis and have been reported to play an essential role in inflammatory symptoms associated with COVID-19 (Lee *et al*, 2023; Zhang *et al*, 2020). Consistently, the ETV/mPSL combination therapy controlled the host inflammatory response to infection. At 6 dpi, hamsters in the mPSL monotherapy group exhibited high expression of cytokines, chemokines, and ISGs (Fig. 5B), corresponding to the delay in viral clearance (Fig. 2G, H). These results indicate that ETV/mPSL combination therapy controls both viral growth and host inflammatory responses to SARS-CoV-2 infection, leading to better outcomes for recipient animals with COVID-19.

**Fig. 4.**
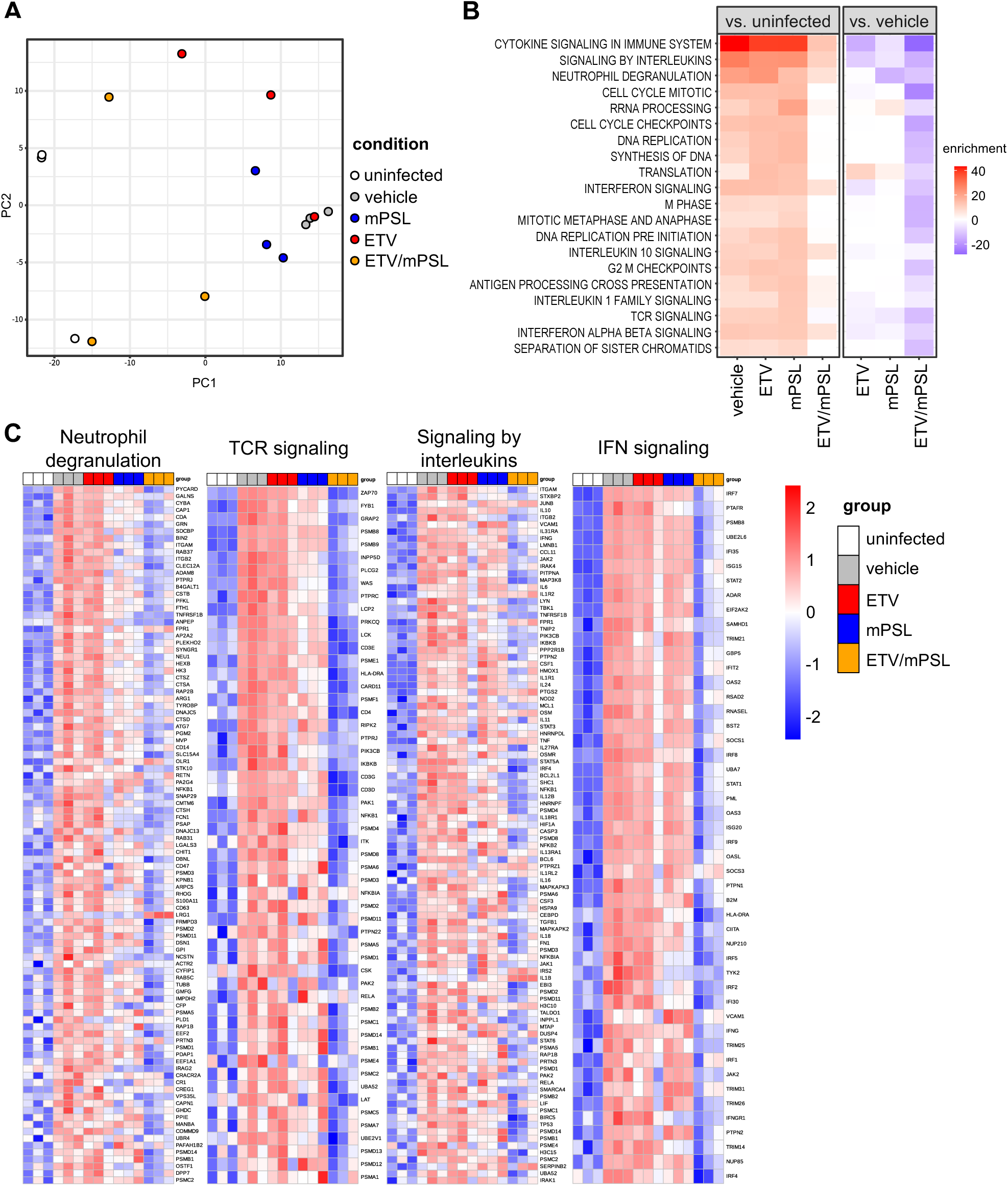
Transcriptomic signatures in the lungs of SARS-CoV-2-infected hamsters receiving therapeutic treatment. Hamsters were infected with 1 × 10^4^ pfu of the SARS-CoV-2 delta variant and treated b.i.d. with ETV (200 mg/kg) and/or mPSL (1 mg/kg) at 2–3 dpi following the schedule shown in Fig. 2A. **A**, Principal component analysis of gene expression data derived from the whole lungs of hamsters at 4 dpi. Dots indicate infected hamsters receiving vehicle (gray), mPSL (blue), ETV (red), or ETV/mPSL (orange) and uninfected controls (white). The explained variance by each component was 65% for PC1 and 18% for PC2. **B,** Gene set enrichment analysis of differentially expressed genes. Upregulated (red) or downregulated (blue) gene pathways compared to uninfected control (left panel) and infected vehicle-treated hamsters (right panel) are shown. Enrichment values were calculated as −log_10_ (adjusted *p*-value), and downregulated pathways are represented with negative values. **C,** Heatmaps showing the relative expression of a panel of genes in enriched pathways. No more than 100 of the top upregulated genes in each pathway in an untreated vehicle control group compared to the uninfected control group are shown. Each column represents a single sample from the vehicle (gray), mPSL (blue), ETV (red), ETV/mPSL (orange) or uninfected control group (white).

**Fig. 5.**
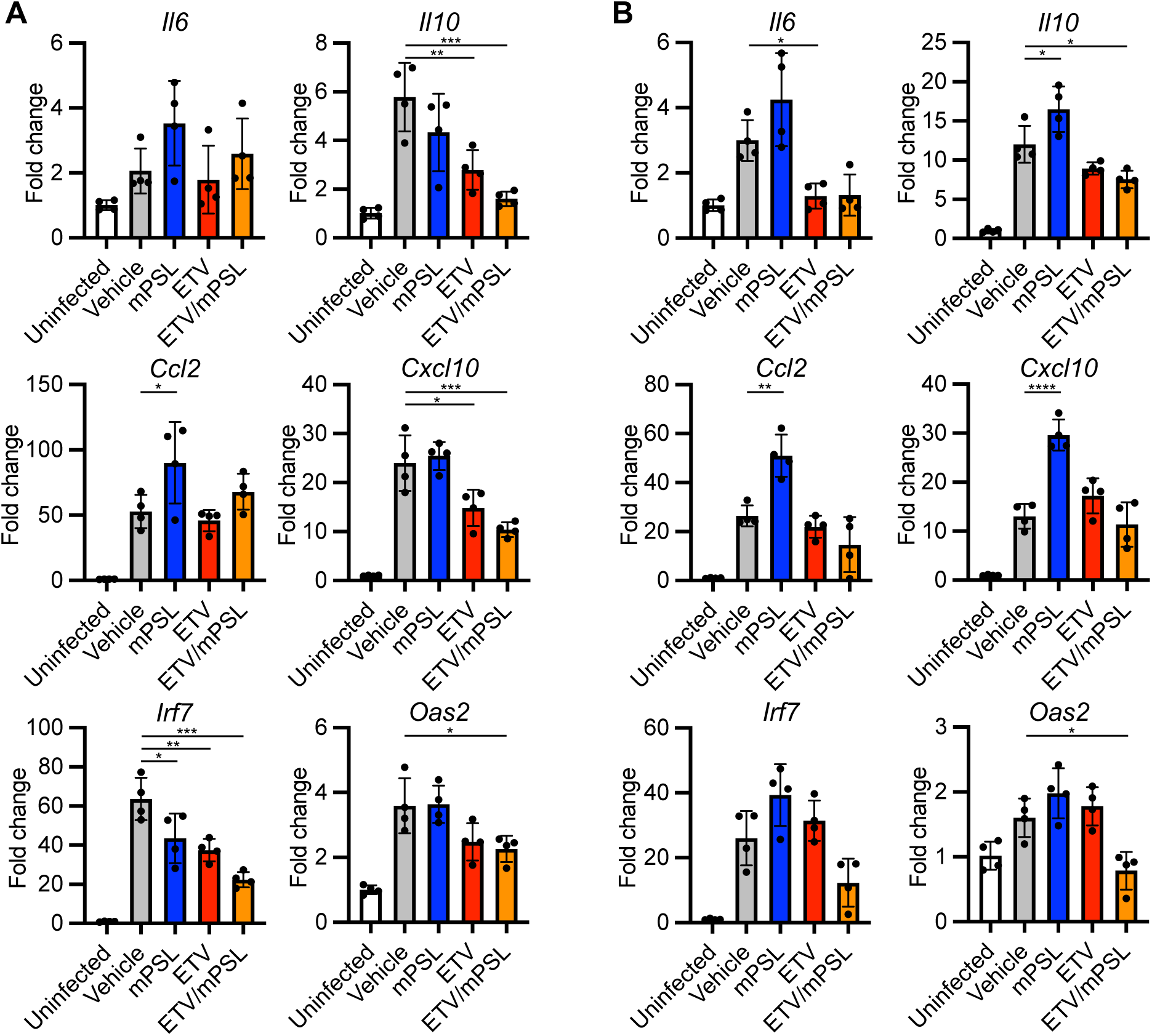
Validation of differential gene expression associated with therapeutic treatment with ETV and mPSL. **A, B**, Relative gene expression of cytokines and interferon-stimulated genes in the lungs of hamsters at 4 (A) and 6 dpi (B). mRNA expression was quantified using a probe-based qRT-PCR assay. Data were normalized to those of *Actb*. The values shown are mean ± SD with each dot representing an individual animal. \**p*<0.05, \*\**p*<0.01, \*\*\**p*<0.001, \*\*\*\**p*<0.0001 by one-way ANOVA with Tukey’s test.

### Combination therapy with ETV and mPSL for SARS-CoV-2 omicron XBB.1

Compared to the highly pathogenic SARS-CoV-2 delta variant, the currently circulating omicron variant has relatively low virulence and slow growth in the lungs, and it causes attenuated disease in hamsters (Halfmann *et al*., 2022; Tamura *et al*, 2023; Yuan *et al*., 2022b). As such, we investigated the effect of ETV/mPSL combination therapy against SARS-CoV-2 XBB.1, a sub-lineage of the omicron variant. Because omicron infection has little impact on body weight and respiratory parameters in hamsters (Tamura *et al*., 2023), we focused on the viral load and inflammation in the lungs to evaluate therapeutic efficacy. ETV monotherapy and ETV/mPSL combination therapy, but not mPSL monotherapy, reduced virus titers and viral RNA loads in the lungs of hamsters at 4 dpi (Fig. 6A, B). Histopathological examination revealed that ETV/mPSL combination therapy improved the severity score with reduced inflammatory cell infiltration and lower numbers of viral RNA/antigen-positive cells (Fig. 6C, D). In addition, mPSL monotherapy attenuated the lung pathology without decreasing the number of virus-infected cells (Fig. 6C, D). The expression of cytokines, chemokines, and ISGs in the lungs was elevated following infection, and this gene upregulation was suppressed by treatment with ETV alone and in combination with mPSL, with these effects presumably related to changes in viral loads (Fig. 6E-J). These results indicate that ETV/mPSL combination therapy ameliorates pneumonia caused by the SARS-CoV-2 omicron variant, whereas ETV and mPSL alone only partially improve host inflammatory responses to the infection.

**Fig. 6.**
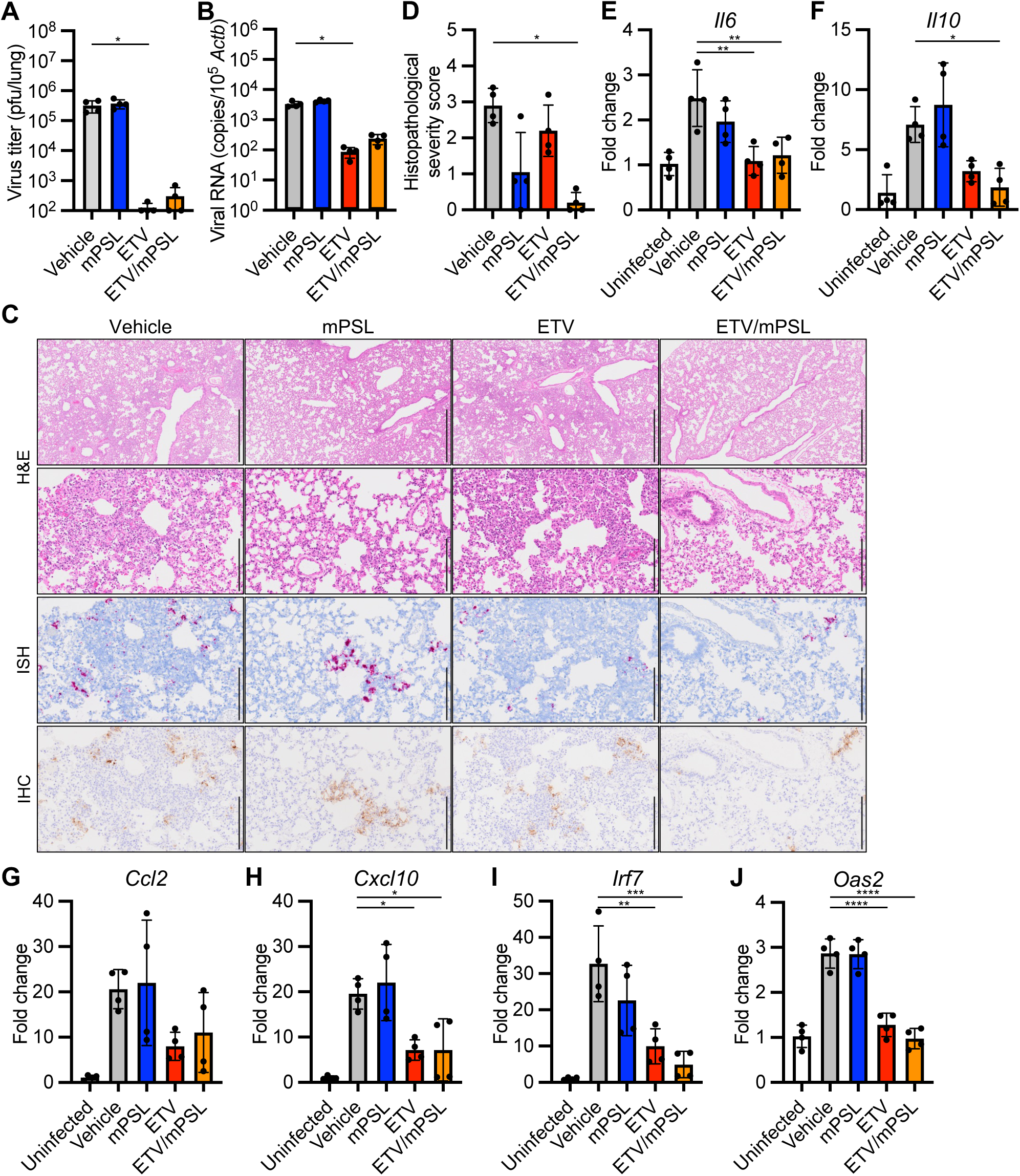
Therapeutic treatment of hamsters infected with SARS-CoV-2 omicron variant. Hamsters were infected with 1 × 10^4^ pfu of the SARS-CoV-2 omicron variant XBB.1 and treated b.i.d. with ETV (200 mg/kg) and/or mPSL (1 mg/kg) at 2–3 dpi. **A**, Virus titers in hamster lungs at 4 dpi were determined by plaque assay. **B**, Viral RNA copies in hamster lungs at 4 dpi were quantified by qRT-PCR and normalized to those of *Actb*. **C**, Representative histopathological images of hamster lungs at 4 dpi. Upper two panels, hematoxylin and eosin staining. The third from the top panels, ISH targeting the nucleocapsid gene of SARS-CoV-2. Bottom panels, IHC targeting the nucleocapsid protein of SARS-CoV-2. Scale bars in the top panels, 1 mm. Scale bars in other panels, 200 μm. **D**, Histopathological severity score of pneumonia based on the percentage area of alveolitis in a given section. **E**–**J**, Relative gene expression of *Il6* (E), *Il10* (F), *Ccl2* (G), *Cxcl10* (H), *Irf7* (I), and *Oas2* (J) in hamster lungs at 4 dpi compared to the findings in the lungs of uninfected hamsters were examined using qRT-PCR. Data were normalized to those of *Actb*. The values shown are mean ± SD with each dot representing an individual animal. \**p*<0.05, \*\**p*<0.01, \*\*\**p*<0.001, \*\*\*\**p*<0.0001 by Kruskal-Wallis test with Dunn’s test (A, B, D) and one-way ANOVA with Tukey’s test (E–J).

## Discussion

Lung bronchial and alveolar epithelial cells are targets for SARS-CoV-2 infection. The infected cells release inflammatory mediators and trigger host inflammatory responses with pulmonary infiltration of immune cells, leading to COVID-19 pneumonia (Lamers & Haagmans, 2022). Therefore, inhibiting SARS-CoV-2 proliferation in the early phase of infection prevents hyperinflammation and mitigates the severity of the disease. In this study, hamsters receiving delayed treatment with ETV from 2 dpi showed greater weight loss, more severe respiratory conditions, and higher expression of cytokines and chemokines than hamsters receiving ETV in an earlier phase of infection, indicating that delayed antiviral treatment resulted in reduced therapeutic efficacy. It is suggested that controlling the host inflammatory response is an essential factor for a better performance of delayed therapy for COVID-19. Because viral loads have been reported to peak around 2 dpi in the lungs of hamsters experimentally inoculated with SARS-CoV-2 (Campos *et al*, 2021; Sia *et al*, 2020), drug administration was initiated in our delayed protocol after substantial viral proliferation, which is associated with the stimulation of host inflammatory responses.

Our mRNA-seq analysis revealed that mPSL monotherapy suppressed neutrophil and T cell activation, suggesting that these phenotypes contribute to the amelioration of COVID-19-induced lung pathology. Consistent with our findings, it has been reported that dexamethasone induced the immunosuppression of neutrophils in patients with COVID-19 (Sinha *et al*, 2022). Conversely, mPSL monotherapy was associated with an increased viral load, delayed viral clearance, and prolonged several types of cytokine/chemokine upregulation in the lungs. These paradoxical effects of steroidal anti-inflammatory drugs were also recognized in a previous study using SARS-CoV-2-infected hamsters (Ye *et al*., 2021). Furthermore, treatment with steroidal anti-inflammatory drugs had varied and inconsistent effects on the expression of cytokine and chemokine genes in various studies, presumably because of differences in the time course, administration route, and doses of the anti-inflammatory drugs (Wyler *et al*., 2022; Ye *et al*., 2021; Yuan *et al*., 2022a). Our histopathological examination revealed that mPSL monotherapy partially improved the lung pathology in hamsters infected with the omicron variant, but not in those infected with the highly pathogenic delta variant. These results highlight the complexity and difficulty of treatment with steroidal anti-inflammatory drugs for COVID-19.

Theoretically, the antiviral effects of drugs could compensate for the undesirable delay in viral clearance induced by treatment with anti-inflammatory drugs in COVID-19. Compared to the effects of ETV or mPSL alone, the combination therapy controlled a wide range of host inflammatory responses, improved the lung pathology, and ameliorated clinical aspects of COVID-19 in hamsters. Furthermore, the combination treatment was effective in infection with both highly pathogenic delta and circulating omicron variants. We propose a model in which ETV/mPSL combination treatment suppressed pulmonary inflammation without delay of virus clearance and achieved strong therapeutic efficacy following delayed treatment in SARS-CoV-2-infected hamsters (Fig. S2). In conclusion, our study demonstrates that combination treatment with antiviral and anti-inflammatory drugs could provide a potent treatment option for COVID-19.

On the other hands, we note some limitations in this study. First, we examined the performance of combination therapy using a single set of oral antiviral/anti-inflammatory drugs. Nirmatrelvir and molnupiravir have been widely used as oral antiviral drugs for COVID-19, and they are candidates for further investigation of antiviral/anti-inflammatory combination therapy. Second, we did not consider the effect of drug–drug interactions between ETV and mPSL in hamsters receiving this combination therapy. A clinical study reported that ETV increases the area under the plasma concentration-time curve of dexamethasone, but ETV has no clinically meaningful effect on the pharmacokinetics of prednisolone in humans (Shimizu *et al*, 2023). We therefore did not include dexamethasone in the combination regimen in this study. Third, this study was conducted using hamsters experimentally inoculated with SARS-CoV-2. Although hamsters mimic many aspects of human COVID-19, experimental settings including the virus strain, virus titer, and inoculum size alter the course and severity of COVID-19 (Handley *et al*, 2023). Most of our studies were conducted using the highly pathogenic delta variant with a large inoculum volume, leading to acute and severe infection. This severe COVID-19 model has an advantage of low individual variation, but it would be less sensitive to observe other therapeutic treatments.

In summary, our study revealed the advantage of combination therapy with ETV and mPSL from the perspective of lung pathology and host inflammatory responses. Since patients can easily receive oral medications, combination therapy with oral antiviral and anti-inflammatory drugs could be a practical and potent treatment option for COVID-19.

## Materials and Methods

### Cells

Vero-TMPRSS2 cells [Vero E6 cells (ATCC, CRL-1586) stably expressing human TMPRSS2] and Vero-hACE2-TMPRSS2 (Vero E6 cells stably expressing human ACE2 and human TMPRSS2) were established as previously described (Sasaki *et al*, 2021b; Uemura *et al*, 2021) and maintained in Dulbecco’s Modified Eagle’s Medium (DMEM) containing 10% fetal bovine serum (FBS).

### Viruses

SARS-CoV-2 delta (strain TY11-927, lineage AY.122, GISAID: EPI_ISL_2158617) and omicron (strain TY41-795, lineage XBB.1, GISAID: EPI_ISL_ 15669344) variants were obtained from the National Institute of Infectious Diseases, Japan. The working viral stocks were prepared by passage on Vero-TMPRSS2 or Vero-hACE2-TMPRSS2 cells. The titers of the prepared virus stocks were determined as plaque forming unit per ml (pfu/ml) using a plaque assay.

### Plaque assay

Vero-TMPRSS2 and Vero-hACE2-TMPRSS2 cells were used for titration of delta and omicron variants, respectively. Cells were inoculated with serial dilutions of either virus stock or clarified tissue homogenates for 1h at 37°C. The cells were then overlaid with DMEM containing 2% FBS, 0.5% Bacto Agar (Becton Dickinson) and 25 μg/ml gentamicin (Wako). At 2 dpi with delta variant and 3 dpi with omicron variant, cells were fixed with 3.7% formaldehyde in phosphate-buffered saline (PBS) and stained with 1% crystal violet.

### Compounds

ETV (also known as code S-217622, fumaric acid co-crystal form) was provided from Shionogi & Co., Ltd.(Unoh *et al*, 2022). mPSL was obtained from the Tokyo Chemical Industry.

### Experimental infection of hamsters

The animal experiments with virus infection were performed in accordance with the National University Corporation, Hokkaido University Regulations on Animal Experimentation. The protocol was reviewed and approved by the Institutional Animal Care and Use Committee of Hokkaido University (approval no. 20-0060). Five-weeks-old male Syrian hamsters (Japan SLC) were intranasally inoculated with 1 × 10^4^ pfu of SARS-CoV-2 in 200 μl of PBS under anesthesia with isoflurane inhalation. ETV and mPSL were suspended in 0.5 % (w/v) methyl cellulose 400. Hamsters were orally administered twice daily with ETV and/or mPSL for 5 days from 0, 1 or 2 dpi under anesthesia with isoflurane inhalation. Vehicle control hamsters were administered with 0.5 % (w/v) methyl cellulose 400. A subset of hamsters at 4 dpi or 6 dpi were sacrificed for lung collection. The lung tissues were used for virus titration, gene expression assays and histopathological examination. Another subset of hamsters were monitored for up to 10 dpi for body weight change. Body weights of animals were monitored daily.

### Virus neutralization assay

To determine the titers of neutralizing antibody in hamsters, serum samples were collected from hamsters at 18 dpi and heat-inactivated at 56°C for 30 min. Virus neutralization assays were performed following the protocol as previously described (Sasaki *et al*, 2021a). The neutralization titer was defined as the reciprocal of the highest serum dilution that completely inhibited the virus cytopathic effect in Vero-TMPRSS2 cells.

### Virus titration and quantitative RT-PCR (qRT-PCR) assays

Lungs from hamsters at 4 or 6 dpi were homogenized in PBS with TissueRuptor (Qiagen). A part of the whole lung homogenate was subjected to plaque assays for virus titration as described above. For measurement of host gene expressions and viral RNA levels, total RNA was extracted from the homogenate with a combination of TRIzol LS (Invitrogen) and Direct-zol RNA MiniPrep kit (Zymo Research). RNA samples were analyzed by qRT-PCR with Thunderbird Probe One-step qRT-PCR Kit (Toyobo) and QuantStudio 7 Flex Real-time PCR system (Applied Biosystems; Thermo Fisher Scientific). Target RNA levels were normalized to hamster *Actb* and calculated by the relative standard curve method. The copy numbers of hamster *Actb* and SARS-CoV-2 nucleocapsid genes were also estimated by the standard curve method. Primers and probes for hamster *Irf7* and *Oas2* were obtained as a PrimeTime Predesigned qPCR assay (Integrated DNA Technologies). Other primers and probes for hamster *Actb*, *Il6*, *Il10*, *Ccl2*, *Cxcl10,* and SARS-CoV-2 nucleocapsid genes were as previously described (Bricker *et al*, 2021; Shirato *et al*, 2020; Zivcec *et al*, 2011).

### Differentially expressed genes

For mRNA-seq analysis, total RNA samples from lungs at 4 dpi were subjected to library preparation using NEBNext Ultra II Directional RNA Library prep kit for Illumina (New England Biolabs) and sequenced on an Illumina NovaSeq 6000 with 150 base pair-end reads at Azenta. Raw mRNA-seq data were deposited to DDBJ with accession IDs DRR477693-DRR477707. Obtained mRNA-seq reads were processed with the nf-core/RNAseq pipeline (Ewels *et al*, 2020). Preprocessed paired reads were aligned to reference genomes with STAR (v2.7.10) (Dobin *et al*, 2013). Genome data for Syrian Golden hamster (Ensemble MesAur1.0) and SARS-CoV-2 (Ensemble ASM985889v3 Wuhan-Hu-1 isolate), respectively, were used (Cunningham *et al*, 2022). Gene level transcript abundances were estimated by Salmon (v1.9.0) (Patro *et al*, 2017) and tximport (v1.22) (Soneson *et al*, 2015). Reads mapped to SARS-CoV-2 genome were removed in further analysis. Differentially expressed gene analysis was performed using DESeq2 (v1.34) (Love *et al*, 2014). Differentially expressed genes were ranked with a *p*-value and the direction of fold change as described elsewhere (Van den Berge *et al*, 2018) and ranked gene lists were used for geneset enrichment analysis (Mootha *et al*, 2003) using ClusterProfiler (v4.2.2) (Wu *et al*, 2021). To utilize human genesets, hamster gene IDs were converted to human gene IDs by g:Profiler suite (Raudvere *et al*, 2019). Genes with one to one ortholog relation were used. Human genesets retrieved from Reactome Pathway (Gillespie *et al*, 2022) and CellMarker (Zhang *et al*, 2019) were used. Cell type markers with evidence code “Experimental” were used. For heatmap analysis, a DESeq2 normalized gene expression table was log2 transformed and was scaled in gene-wise manner to establish relative gene expression values (z-score).

### Pulmonary function tests

Pulmonary function was assessed using a whole-body plethysmography system (Data Sciences International) as previously described (Yamasoba *et al*, 2022). In brief, hamsters were placed individually in unrestrained plethysmography chambers. After 30 sec for acclimation, respiratory parameters were acquired over a 3-min period by using FinePointe software (Data Sciences International).

### Histopathological examination

Lung tissues at 4 dpi were fixed in 3.7% formaldehyde in PBS and embedded in paraffin. To detect viral RNA in the paraffin sections, *in situ* hybridization was carried out using an RNA scope 2.5 HD Red Detection kit (Advanced Cell Diagnostics) with an antisense probe targeting the nucleocapsid gene of SARS-CoV-2 (Advanced Cell Diagnostics) as previously described (Halfmann *et al*., 2022). To detect viral antigen in the sections, IHC was carried out using a rabbit monoclonal antibody for SARS-CoV nucleocapsid protein (catalogue #40143-R001, clone #001, Sino Biological) which cross-reacts with SARS-CoV-2 nucleocapsid protein. Specific antigen-antibody reactions were visualized by means of 3,3’-diaminobenzidine tetrahydrochloride staining using the Dako Envision system (Dako Cytomation). Histopathological severity score of pneumonia was determined based on the percentage of alveolar inflammation in a given area of a pulmonary section collected from each animal in each group using the following scoring system: 0, no inflammation; 1, affected area (≤1%); 2, affected area (>1%, ≤10%); 3, affected area (>10%, ≤50%); 4, affected area (>50%); an additional point was added when pulmonary edema and/or alveolar hemorrhage was observed (Uraki *et al*, 2022a). The average score for all lobes was calculated for each animal.

### Statistical analysis

Statistical significance was determined by two-way analysis of variance (ANOVA) with Dunnett’s test (Figs. 1B and 2B–D), one-way ANOVA with Tukey’s test (Figs. 1C–D, 1H–K, 5A–B, 6E–J) and Kruskal-Wallis test with the Dunn’s multiple comparisons test (Figs. 1E–F, 2E–H, 3C–D, 6A–B, 6D). All statistical tests were carried out using Prism version 9.5.1 (GraphPad software).

## Acknowledgments

We thank Dr. Ken Maeda in the National Institute of Infectious Diseases, Japan for providing the SARS-CoV-2 variants and Dr. Kei Sato in Tokyo University, Japan for supporting pulmonary function tests using a plethysmography system. We also thank Yuko Sato and Seiya Ozono for technical assistance. This work was supported by the Japan Agency for Medical Research and Development (AMED) under Grant numbers JP23wm0125008, JP223fa627005 and JP23fk0108637; Japan Science and Technology Agency (JST) Moonshot R&D under Grant numbers JPMJMS2025; and the World-leading Innovative and Smart Education (WISE) Program from the Ministry of Education, Culture, Sports, Science, and Technology (MEXT), Japan under Grant numbers 1801.

---

**Fig. S1.**
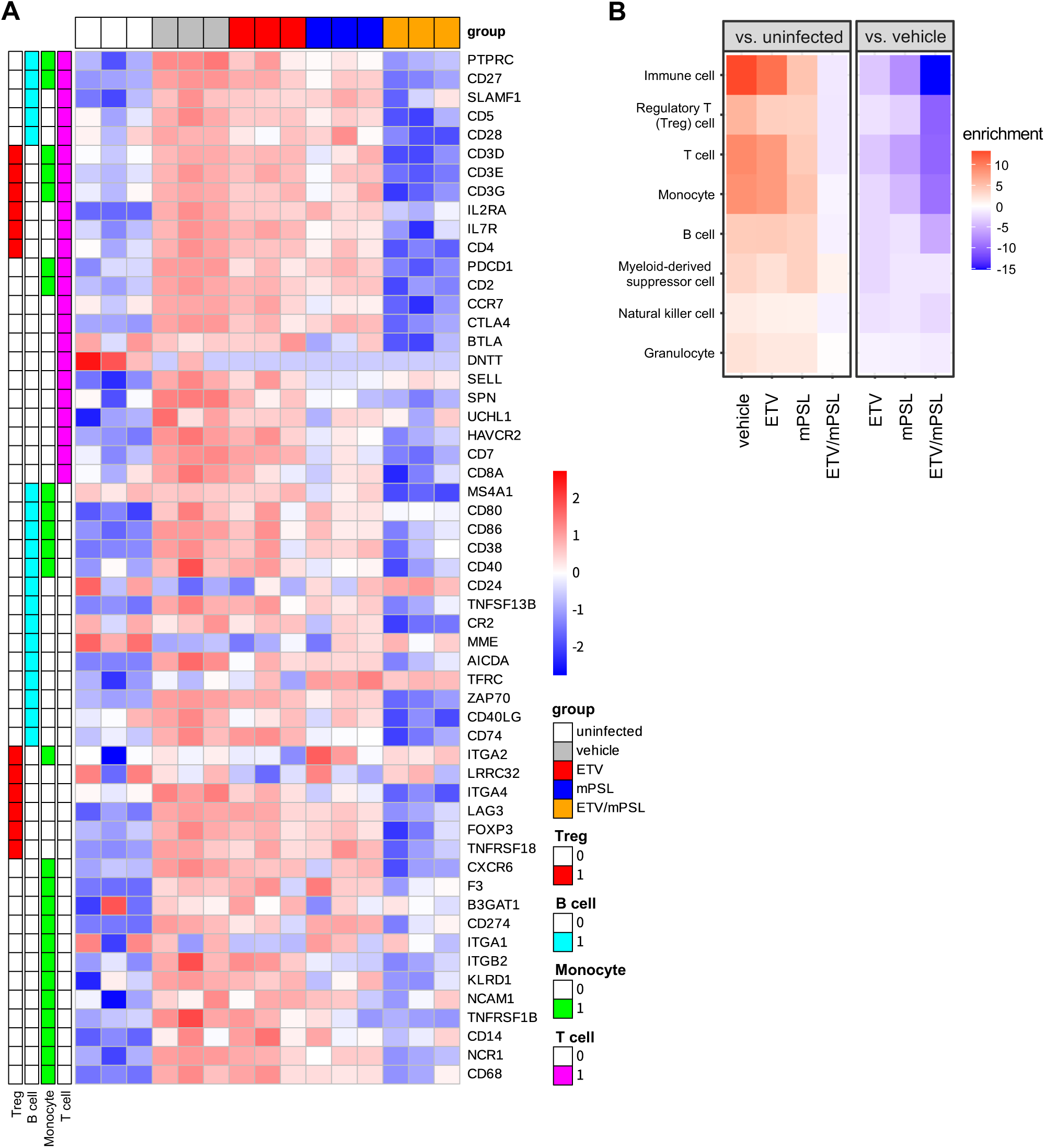
Immune cell signatures in the lungs of SARS-CoV-2-infected hamsters receiving treatment. **A**, Heatmap of gene expression data derived from the whole lungs of hamsters at 4 days post-infection (dpi) as described in Fig. 4 showing the relative gene expression of a panel of immune cell markers in a red (high) to blue (low) color scheme. Each column represents a single sample from the vehicle (gray), methylprednisone (mPSL, blue), ensitrelvir (ETV, red), ETV/mPSL (orange), or uninfected control group (white). Each row represents a marker from the assigned cell types: regulatory T cell (red), B cell (cyan), monocyte (green), and T cell (pink). **B,** Gene set enrichment analysis of canonical cell markers shown in **A**. Upregulated (red) or downregulated (blue) gene pathways compared to the findings in the uninfected control group (left panel) or vehicle-treated group (right panel) are shown. Enrichment values were calculated as −log_10_ (adjusted *p*-value), and downregulated pathways are represented as negative values.

**Fig. S2.**
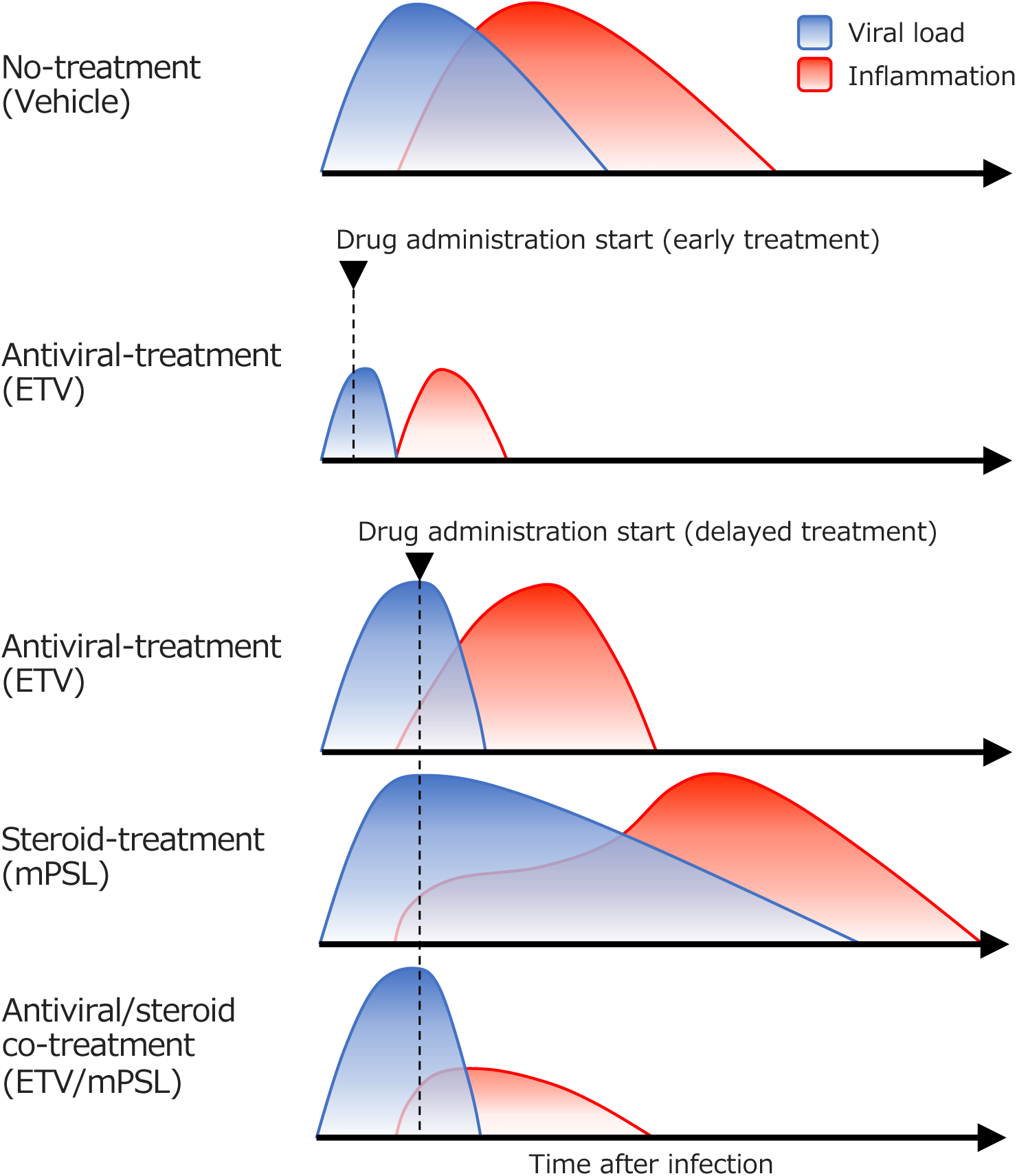
Schematic representation of a proposed model of monotherapy and combination therapy with ETV and mPSL for severe COVID-19. Extensive inflammation is caused by SARS-CoV-2 infection in severe COVID-19. Early treatment with ETV, reduces peak virus titer and prevents the progression of COVID-19. In delayed dosing therapy, ETV monotherapy reduces viral loads, accelerates virus clearance and shortens the inflammatory state. Immunosuppression induced by mPSL monotherapy ameliorates inflammation but hinders viral clearance leading to a prolonged inflammatory state. ETV/mPSL combination therapy controls both viral loads and inflammation resulting in a high therapeutic performance.

